# Partitioning of nuclear material into apoptotic fragments through establishment of asymmetric cell death morphology

**DOI:** 10.64898/2026.07.13.738122

**Authors:** Jascinta P. Santavanond, Lanzhou Jiang, Amy L. Hodge, Dilara C. Ozkocak, Alper Ceviker, Satoko Arakawa, Shigeomi Shimizu, Ikuyo Yoshino, Stephanie F. Rutter, Thanh Kha Phan, Rochelle Tixeira, Amy A. Baxter, Sarah Caruso, Lucas M. Newton, Rebecca Stephens, Patrick O. Humbert, Mark D. Hulett, Georgia K. Atkin-Smith, Ivan K. H. Poon

**Affiliations:** Department of Biochemistry and Chemistry, La Trobe Institute for Molecular Science, La Trobe University, Melbourne, Victoria 3086, Australia; Research Centre for Extracellular Vesicles, La Trobe University, Victoria, Australia; Department of Pathological Cell Biology, Medical Research Institute, Tokyo Medical and Dental University, 1-5-45 Yushima, Bunkyo-ku, Tokyo 113-8510, Japan; VIB-UGent Center for Inflammation Research, Department of Biomedical Molecular Biology, Ghent University, Ghent, Belgium; The Walter and Eliza Hall Institute of Medial Research, Parkville, Victoria, Australia; University of Melbourne, Victoria, Australia

**Keywords:** Apoptosis, apoptotic bodies, cell clearance, DNA, extracellular vesicles, nuclear material

## Abstract

Cellular material in apoptotic cells must be efficiently cleared by phagocytes to maintain tissue homeostasis. Defects in this process can lead to the onset of secondary necrosis and the release of intracellular contents such as damage associated molecular patterns (DAMPs) and autoantigens that are often derived from the nucleus. Therefore, appropriate handling and clearance of apoptotic material is vital to prevent unwanted inflammatory response and the onset of autoimmune disorders. However, how nuclear material is packaged by apoptotic cells for effective clearance by phagocytes is not well understood. By utilising murine models of apoptosis, we observed that a distinct subset of large extracellular vesicles generated from apoptotic thymocytes, known as apoptotic bodies (ApoBDs), can harbour the majority of nuclear contents. Mechanistically, we discovered that apoptotic cells can asymmetrically partition the nucleus into a single large membrane bleb located at one side of the cell, with other cellular contents such as mitochondria and acid organelles distributed to the opposite side. Whilst this newly observed apoptotic morphology, coined as asymmetric cell death morphology (AsyCDM), is morphologically similar to the process of erythroblast enucleation, pharmacological compounds that could interfere with erythroblast enucleation did not block the establishment of AsyCDM during apoptosis. Notably, AsyCDM was reliant on the contractile forces generated by ROCK1-dependent plasma membrane blebbing. Taken together, this study suggests that intracellular contents are partitioned into different ApoBD subsets during apoptosis through a regulated process driven by ROCK1-dependent actomyosin contraction.

## Introduction

Billions of cells undergo cell death daily in the human body and cell corpses are cleared rapidly to ensure cellular contents are removed and recycled. Amongst the various cell death modalities, apoptosis is the most frequently studied. If apoptotic cells are not removed efficiently by phagocytes through the process known as efferocytosis, the integrity of the plasma membrane can become compromised and damage associated molecular patterns (DAMPs) such as ATP and heat shock proteins (HSPs) are released, subsequently eliciting inflammatory responses^1–3^. Notably, the nucleus can harbour many DAMPs and autoantigens including DNA, high mobility group box 1 (HMGB1) and histones. Thus, appropriate disposal of nuclear material from dying cells has been suggested to be key in preventing unwanted inflammation and development of autoimmunity^4^. For example, mice lacking the key serum nuclease DNase I develop systemic lupus erythematosus-like disease, indicated by the presence of anti-nuclear antibodies in the serum and immune-complex glomerulonephritis^5^.

As the nucleus is one of the largest organelles within the cell, a number of mechanisms work in concert to ensure nuclear material within apoptotic cells are disposed of effectively. Notably, degradation of apoptotic cell-derived DNA occurs in two stages. Activation of caspase 3 during apoptosis promotes the cleavage of inhibitor of caspase-activated DNase (ICAD) in the cytoplasm and subsequently enables caspase-activated DNase (CAD) to enter the nucleus to mediate degradation of chromosomal DNA^6,7^. Engulfment of DNA-containing apoptotic material by macrophages can further aid the degradation of chromosomal DNA via DNase II within phagolysosomes^4,8^. Concurrently during apoptosis, the structural integrity of the nucleus is destabilised by the proteolysis of proteins such as lamin A/C and lamin-associated proteins^9^, as well as membrane blebbing, leading to nuclear fragmentation^10,11^. Such destabilisation of nuclear contents during apoptosis is thought to aid their processing within phagocytic cells^12^.

During apoptosis, dying cells can undergo distinct morphological changes including the formation of membrane blebs and thin membrane protrusion to aid cell disassembly into membrane-bound fragments called apoptotic bodies (ApoBDs) that are approximately 1-5 μm in diameter^13^. It has been well-described that organelles such as the nucleus and mitochondria can be distributed into ApoBDs^14–17^. However, whether nuclear material are packaged into distinct ApoBDs for cell clearance and whether the distribution of cellular contents during apoptosis is regulated is not well understood. In this study, we describe the establishment of a novel asymmetric morphology that can aid the partitioning of nuclear material during apoptosis. Particularly, the nucleus and various cytoplasmic organelles such as mitochondria are distributed asymmetrically in apoptotic cells during this asymmetric morphological stage and subsequently aid the formation of distinct ApoBDs that harbour the majority of nuclear contents. Mechanistically, ROCK1-mediated actomyosin contraction is required for the establishment of asymmetric cell morphology and partitioning of nuclear contents during apoptosis.

## Materials and Methods

### Reagents

Trovafloxacin, Hoechst 33342, PKH67, PKH26, NucRed, SYTOX Green, KN-93, EDTA, monensin sodium salt, anti-Fas (clone CH11) and raptinal were purchased from Sigma-Aldrich. MitoTracker Green, TO-PRO-3, anti-mouse CD4 PE-Cy7 (GK1.5), anti-mouse CD8α PE (53-6.7), anti-mouse CaMK-II (6G9), Donkey anti-mouse IgG (H+L) secondary antibody Alexa Fluor 488 and Donkey anti-mouse IgG (H+L) secondary antibody Alexa Fluor 568 were purchased from Thermo-Fisher Scientific. Anti-mouse Ter119, anti-mouse Ter119 Alexa Fluor 647 and anti-mouse/human CD44 PE/Cy7 (IM7) were purchased from BioLegend. GSK 269962 was purchased from Tocris Bioscience. ABT-737 and S63845 were purchased from Jomar Life Research. Golgi ID-Green was purchased from Enzo life sciences. A5 FITC, A5 PE and 10 × A5 binding buffer were purchased from BD Biosciences. SiR-Actin and SiR-Tubulin live cell probes were purchased from Spirochrome. ABT-737 and S63845 were purchased from Jomar Life Research. SPHERO^TM^ AccuCount Blank Particles (5.0-5.9 μm) was purchased from Spherotech.

### *In vivo* model of apoptosis

5-week-old C57bl/6 mice (male and female) were maintained and used under approval of AEC21034, La Trobe University. All methods and experiments were approved by the La Trobe University Animal Ethics Committee in accordance with the National Health & Medical Research Council Australia (NHMRC) code of practice for the care and use of animals for scientific purposes. Mice were intraperitoneally injected with 12.5 mg/kg dexamethasone (dex) to induce apoptosis *in vivo* as previously described^15^. Whole-body x-ray irradiation treatment was performed using the Rad Source RS-2000 X-Ray Irradiator (Rad Source Technologies Inc). Mice were housed in an Allentown cage and exposed to 6.8 Gy. Following treatment, mice were culled after 6 h by CO_2_ and the thymus was harvested to generate single cell suspensions for flow cytometry analysis as described below.

### Cell culture

Human THP-1 monocytic, Jurkat T (clone E6-1), A549 epithelial cells and HeLa epithelial cells were obtained from ATCC (TIB-202, TIB-152, CCL-185 and CCL-2, respectively). A431 epithelial cells were obtained from Lonza. DC2.4 DC-like cells were a kind gift from Dr Kenneth L Rock, University of Massachusetts Medical School. PANX1^-/-^ and ROCK1^-/-^ GFP^+^ mCherry^+^ (clonal) Jurkat T cells were generated by a CRISPR/Cas9-based approach^18^. A431 H2B-GFP cells were generated by transfecting A431 epithelial cells with H2B-GFP (a kind gift from Geoff Wahl, Addgene plasmid #11680; http://n2t.net/addgene:11680; RRID:Addgene_11680) using Lipofectamine 2000 (Thermo-Fisher Scientific) according to manufacturer’s protocol. All cell lines were cultured in RPMI 1640 (Gibco) or MEM (Lonza) medium containing 10% (v/v) fetal bovine serum (Scientifix), penicillin (50 U/mL), streptomycin (50 μg/mL) and 0.2% MycoZap (Lonza), at 37°C in 5% CO_2_. MCF10A Cas9 and Scribble^-/-^ cells were generated by CRISPR/Cas9-based approach^19^ and were maintained in Dulbecco’s modified Eagle’s medium with F12 (DMEM:F12; Gibco), supplemented with 5% (v/v) horse serum (Gibco), 10μg/ml insulin (Novo Nordisk), 0.5μg/ml hydrocortisone (Sigma-Aldrich), 20ng/ml human epidermal growth factor (Sigma-Aldrich), 100ng/ml cholera toxin (List Biological Labs), 100ng/ml penicillin/streptomycin, 2mM glutamine at 37°C in 5% CO_2_. Orthochromatic erythroblasts were isolated from mouse spleen by fluorescence activated cell sorting using FACS Aria Fusion (BD Bioscience) as previously described^20^. Ter119 and CD44 positive cells were collected and seeded at a density of 1.5 x10^5^ cells/mL in MEM-α supplemented with 10% (v/v) foetal bovine serum, 1 mM sodium pyruvate and 1x GlutaMAX. Cells were incubated at 37°C in 5% CO_2_ for a minimum 5 h to allow for enucleation.

### Immunoblotting

Whole cell samples were lysed in lysis buffer (1% IGEPAL® CA-630, 10% glycerol, 1% Triton-X-100, 150 mM NaCl, 20 mM HEPES pH 7.4, protease inhibitor cocktail tablet (Roche)) and were analysed by SDS-PAGE and immunoblotting using the following antibodies: anti-Scribble monoclonal antibody (7C6.D10; 1:100; Sigma-Aldrich), anti-Erk2 (12A4, 1:1,000; Santa Cruz) and HRP-conjugated sheep anti-mouse Ig (1:5,000; GE Healthcare).

### Preparation of human peripheral blood mononuclear cells (PBMC), isolation of PBMC-derived monocytes and generation of monocyte-derived dendritic cells

Fresh buffy coat was obtained from the Australia Red Cross Blood Service (Melbourne, Australia) with ethics approval (FHEC09/16) from La Trobe Human Ethics Committee. Human PBMCs and PBMC-derived monocytes were isolated as per previous study^16^. For the generation of immature dendritic cell (iDC), monocytes were cultured in complete RPMI containing 10% FCS, 1,000 U/mL human GM-CSF (Miltenyi Biotech), and 400 U/mL human IL-4 (Miltenyi Biotech). Cells were cultured in complete medium, and iDCs (cells in suspension) were used on the seventh day.

### Induction of cell death *in vitro*

Cells were either exposed to ultraviolet (UV) irradiation at 150 mJ/cm^2^ using the Stratagene UV stratalinker 1800 (Agilent Technologies), treated with anti-Fas (250 ng/mL), BH3 mimetic cocktail ABT-737/S63845 (2.5 μM/0.5 μM) or raptinal (5 μM) to induce apoptosis. After UV or anti-Fas treatment, cells were incubated at 37°C, 5% CO_2_ for 4 h. Primary thymocytes (collected from 4- to 5-week-old C57BL/6 mice, female) were treated with 50 μM dex for 6 h. During apoptosis induction, all cell types were incubated in 1% BSA/RPMI media. To induce influenza A virus (IAV) mediated cell death, A431 epithelial cells were infected with WT-PR8 (A/Puerto Rico/8/1934 H1N1) at a multiplicity of infection of 10 in acidified medium for 1 h and incubated for 24 h. To induce necrosis by hyperthermic treatment, Jurkat T cells were incubated in a 56°C water bath for 1 h in 1% BSA/RPMI media.

### Flow cytometry analysis

Thymocyte single cell suspensions were prepared by mechanical digestion of thymus in complete RPMI media supplemented with 1 mg/mL collagenase type 1 (Thermo-Fisher Scientific). Samples were resuspended at room temperature for 20 min. EDTA pH 8.0 was added to a final concentration of 12 mM and cells were resuspended for a further 5 min on ice. Cell suspensions were filtered through a 70 μm cell strainer, pelleted at 3,000 x *g* then resuspended in complete RPMI media. For flow cytometry analysis, cell suspensions were stain with anti-mouse CD4 PE-Cy7 (1:300), anti-mouse CD8α PE (1:300), A5 FITC (1:100), TO-PRO-3 (1:2,000) and Hoechst 33342 (1:2,000) diluted in 1x A5 binding buffer at room temperature for 10 min. For flow cytometry analysis of Jurkat T cells, cells were induced to undergo apoptosis as described above then stained with A5 FITC (1:200), TO-PRO-3 (1:2,000) and Hoechst 33342 (1:2,000) diluted in 1x A5 binding buffer at room temperature for 10 min. All samples were analysed using the BD FACSCanto II flow cytometer (BD Bioscience). Data analysis was performed using FlowJo software 10.10 (BD Bioscience).

### Histological analysis

Whole thymic tissue was harvested and fixed in 4% paraformaldehyde (PFA) for 24 h and processed using the Tissue-Tek VIP 6 AI Tissue Processor (Sakura Finetek). Antigen retrieval was performed using the EnVision FLEX TRS, high pH (Dako) at 97°C for 30 mins. Paraffin-embedded sections were immunohistochemically stained with anti-F4/80 (clone D2S9R, 1:500, Cell Signalling Technology) and counterstained with Mayer Hematoxylin using the automated Omnis EnVision G2 template (Dako, Glostrup) as previously described^21^.

### Differential interference contrast confocal microscopy

4-well or 8-well Nunc^TM^ Lab-Tek^TM^ II chambered coverglass (Nunc) were used for live cell imaging. Approximately 1.2×10^5^ Cells were seeded per well of a 4-well chamber slide system (or 6×10^4^ cells/well of an 8-well chamber slide system) in 1% BSA/RPMI medium. For drug treatments, compounds in 1% BSA/RPMI were added to the cells in the chambers. Time-lapse differential interference contrast (DIC) microscopy was performed at 37°C with 5% CO_2_ using Cell Observer spinning disk confocal (Zeiss).

### Monitoring the localisation of organelles and cell surface markers by confocal microscopy

Live cell imaging to detect fluorescent probes was performed with either a Zeiss Confocal Spinning Disk, Zeiss Confocal LSM 900 or the Zeiss Confocal LSM 780 PicoQuant FLIM microscope using ×63 oil immersion. Jurkat T cells, primary iDCs and A431 squamous epithelial cells were stained with MitoTracker Green, LysoTracker Red, Hoechst 33342, Golgi ID-Green, PKH67, PKH26 or SYTOX Green (Miltenyi Biotec), according to manufacturer’s instructions, before imaging or induction of cell death. Enucleating orthohromatic erythroblast were stained as previously described^22^. Briefly, cells were fixed in 4% PFA, permeabilised with PBS-0.3% Triton X-100 and stained with anti-CaMKII (1:100) and anti-Ter119 (1:100) diluted in blocking buffer (PBS-3% BSA/1% donkey serum) overnight at 4°C. Cells were washed with PBS then stained with secondary antibodies donkey anti-mouse Alexa Fluor 488 (1:300), donkey anti-mouse Alexa Fluor 568 (1:300) and 1 μg/mL DAPI diluted in blocking buffer for 2 h at room temperature. Cells were mounted on microscope slides using a cytospin centrifuge and ProLong Gold antifade reagent. In some experiments, cells were stained with TO-PRO-3 and A5 FITC in 1× AV binding buffer to monitor morphological changes during the progression of apoptosis. Confocal microscopy was performed at 37°C in a humidified atmosphere with 5% CO_2_. Image processing and data analysis were performed using the ZEN Blue software (Zeiss) or ImageJ software (National Institutes of Health).

### Monitoring cytoskeletal components by confocal microscopy

The actin filaments and microtubules of Jurkat T cells were stained with SiR-Actin and SiR-Tubulin probe, respectively, according to the manufacturer’s instruction prior to apoptosis induction. Briefly, cells were stained with 100 nM SiR-Actin or SiR-Tubulin and incubated at 37°C in a humidified atmosphere containing 5% CO_2_ for 1 h and imaged by confocal microscopy.

### Transmission electron microscopy

Jurkat T cells and A431 cells were exposed to UV irradiation at 150 mJ/cm^2^ using a CX-2000 UV crosslinker (UVP Inc.) and incubated at 37°C, 5% CO_2_ for 2 or 4.5 h, respectively. Cells were fixed by Karnofsky solution, washed by distilled water, then fixed in an aqueous solution of 1% OsO_4_ (for membrane fixation) and treated with 1% uranyl acetate solution. Fixed cells were embedded in Epon 812, and thin sections were cut and stained with uranyl acetate and lead citrate for observation under a Jeol-1010 electron microscope (Jeol) at 80 kV.

### Scanning electron microscopy

HeLa and A549 cells were seeded onto disks of ACLAR film pre-coated with collagen in 12-well plates (37°C, 5% CO_2_). Following apoptosis induction, cells were then fixed with 2.5% Glutaraldehyde in 0.1 M phosphate buffer for 1 h at room temperature. After fixation, cells were resuspended in PBS and incubated 4°C overnight. The next day, the PBS was aspirated, and the cells were post-fixed in 1% Osmium tetroxide/Sorenson’s phosphate buffe for 1 h at room temperature, then washed 2 x 10 min in distilled water. Cells were dehydrated in a graded ethanol series (20, 50, 70, 90%) for 10 min followed by 100% ethanol, 3 x 10 min. Samples were dried with hexamethyldisilazane and imaged on a Hitachi SU7000 FE SEM (HITACHI) running at 1 kV.

### Statistical analysis

Unless otherwise described, the data are presented as means ± s.e.m., unpaired Student’s two-tailed t-test was applied to determine statistical significance between two groups (control versus a specific treatment). A *P*-value of less than 0.05 was considered statistically significant.

## Results

### Formation of distinct DNA-containing ApoBDs *in vivo* and *in vitro*

Induction of thymocyte apoptosis is a widely used model to study apoptotic and cell clearance processes. Previous studies using electron microscopy, immunohistochemistry and flow cytometry approaches have demonstrated that thymocytes undergo high levels of cell death and form ApoBDs, which are rapidly cleared by residential phagocytes^15,23–26^. To examine the distribution of DNA into ApoBDs *in vivo*, we induced thymocytes to undergo apoptosis by dexamethasone (dex) or whole-body x-ray irradiation (irradiation) and monitored the formation of ApoBDs in the thymus by flow cytometry (Figure 1A). Thymocyte-derived ApoBDs were identified as FSC^low^/CD4^int^/CD8^int^/A5^int^ and the presence of DNA in ApoBDs was determined by Hoechst 33342 (cell-permeable DNA-specific dye) staining (Supplementary Figure 1). Induction of thymocyte apoptosis *in vivo* by dex treatment resulted in the detection of ∼40% of ApoBDs containing DNA (Figure 1B). Notably, DNA^+^ ApoBDs were larger in size than DNA^-^ ApoBDs, as determined by FSC (Figure 1C). Similar to *in vivo* studies, thymocytes undergoing apoptosis following dex treatment *in vitro* also generated a distinct population of DNA^+^ ApoBDs that were consistently larger than DNA^-^ ApoBDs (Figure 1D-E). Furthermore, consistent with the dex-induced apoptosis model, both irradiation and homeostatic apoptosis led to the detection of distinct DNA^+^ ApoBD populations (∼20%) possessing the same characteristics both *in vivo* and *in vitro* (Figure 1F-I, Supplementary Figure 2A-D). Similar to mouse thymocytes, human Jurkat T cells undergoing anti-Fas or UV irradiation induced apoptosis formed DNA^+^ ApoBDs that were consistently larger than DNA^-^ ApoBDs (Supplementary Figure 2E-H). Collectively, this data demonstrates that a distinct population of larger ApoBDs containing DNA are generated under both *in vivo* and *in vitro* conditions.

**Figure 1.**
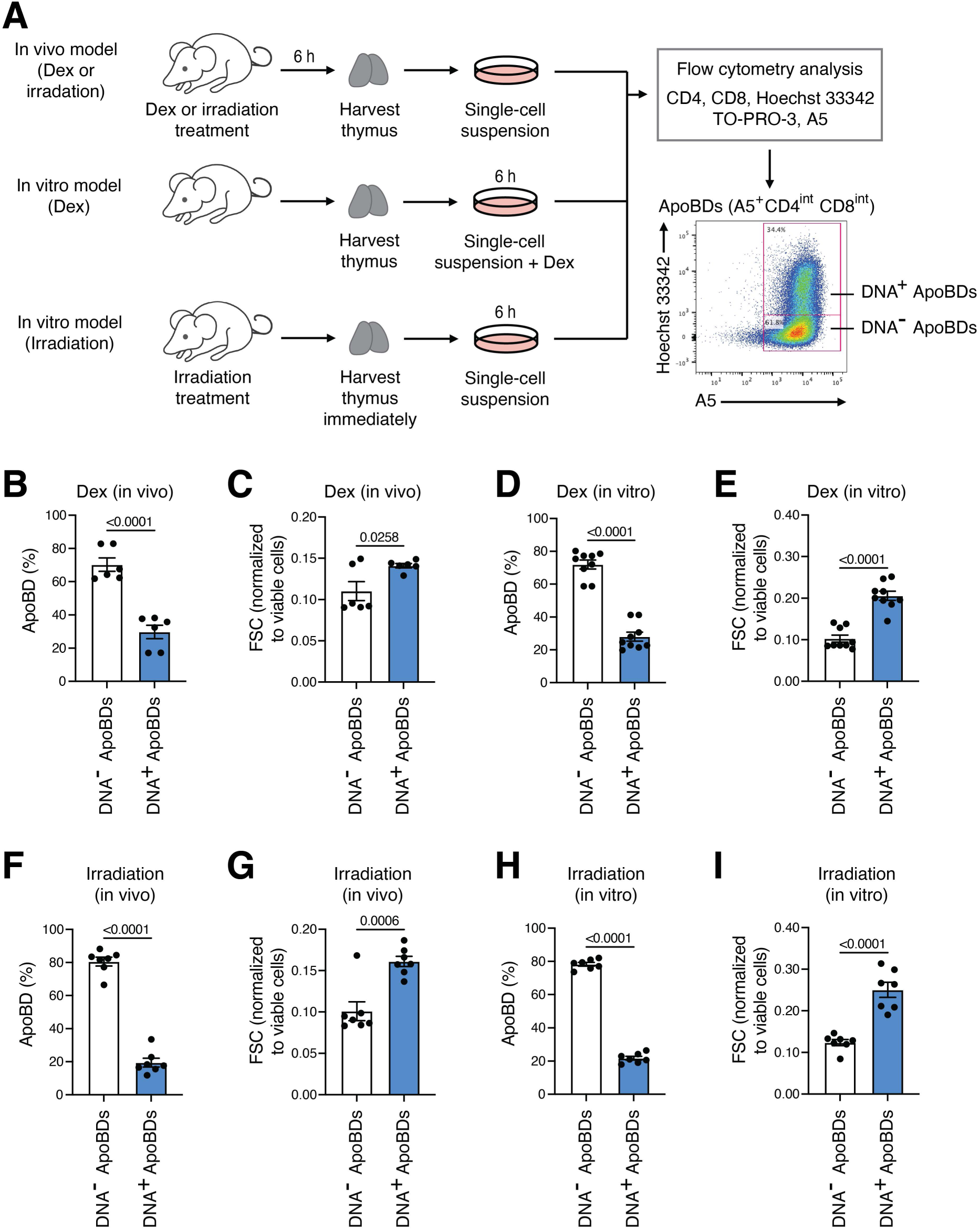
Formation of distinct DNA-containing ApoBDs *in vivo* and *in vitro*. **(A)** Schematic of *in vivo* and *in vitro* dex and irradiation models. **(B)** Flow cytometry analysis of the distribution of DNA into thymocyte-derived ApoBDs following 12.5 mg/kg dexamethasone (dex) treatment *in vivo*. (*n* = 6) Error bars represent s.e.m. **(C)** Relative size of DNA^-^ and DNA^+^ ApoBDs compared to viable cells following 12.5 mg/kg dex treatment *in vivo*. (*n* = 6) Error bars represent s.e.m. **(D)** Flow cytometry analysis of the distribution of DNA into thymocyte-derived ApoBDs following 50 μM dex treatment *in vitro.* (*n* = 9) Error bars represent s.e.m. **(E)** Relative size of DNA^-^ and DNA^+^ ApoBDs compared to viable cells following 50 μM dex treatment *in vitro*. (*n* = 9) Error bars represent s.e.m. Flow cytometry analysis of the distribution of DNA into thymocyte-derived ApoBDs following 6.8 Gy irradiation *in vivo* **(F)** and the relative size compared to viable cells **(G).** (*n* = 7) Error bars represent s.e.m. Flow cytometry analysis of the distribution of DNA into thymocyte-derived ApoBDs following 6.8 Gy irradiation *in vitro* **(H)** and the relative size of DNA^-^ and DNA^+^ ApoBDs compared to viable cells **(I).** (*n* = 7) Error bars represent s.e.m. Unpaired student’s two tailed *t*-test was performed to determine the indicated *p*-value.

### Establishment of asymmetric morphology during apoptosis

Precisely how organelles such as the nucleus are packaged into ApoBDs during apoptosis is not well defined and often thought to be through a stochastic process. To investigate the distribution of various organelles in apoptotic cells, Jurkat T cells and human A431 squamous epithelial cells were stained with Hoechst 33342, MitoTracker Green and LysoTracker Red, to monitor the localisation of the nucleus, mitochondria, and acidic organelles (e.g. lysosome), respectively, by confocal microscopy (Figure 2A). Interestingly, condensed nuclear DNA was distributed into the largest bleb (the single rounded part with a smooth membrane surface) on one side of apoptotic cells, whereas the majority of mitochondria and acidic organelles were localised at the opposite side of the apoptotic cells (Figure 2A). Although for lymphocytes like Jurkat T cells, the nucleus is often positioned to one side in viable cells (Figure 2A), the location of the DNA was frequently found to be located in 1 large bleb at a single extremity of apoptotic cells (Figure 2A and B). We confirmed this asymmetric morphology and distribution of cellular contents by transmission electron microscopy (TEM) analysis of apoptotic Jurkat T cells, A431 cells and mouse DC2.4 dendritic cell (DC)-like cells (Figure 2B). Moreover, scanning electron microscopy (SEM) analysis also provided a high-resolution approach to visualise this distinct asymmetric morphology generated by apoptotic human HeLa cells and A549 epithelial cells treated with BH3 mimetics or infected with influenza A virus, respectively (Figure 2C). As this distinct morphology was not described previously, we coined this as <u>Asy</u>mmetric <u>C</u>ell <u>D</u>eath <u>M</u>orphology (AsyCDM). To quantify the frequency at which cells undergo AsyCDM and partition the nucleus into a single large bleb, we performed time-lapse confocal microscopy to monitored Jurkat T cells undergoing anti-Fas mediated apoptosis. Cells exhibiting AsyCDM were classified into four classes based on the location of the nucleus as monitored by NucRed staining. Class I describes cells that discretely distribute the nucleus into a single large bleb; Class II describes the nucleus in transition between the cell centre and a bleb; Class III describes nuclear fragmentation with nuclear material located at multiple sites; and Class IV describes when the nuclear material remains in the central body of the blebbing cell (Figure 2D, Supplementary Figure 3A). By using this categorisation approach, we observed that cells undergoing AsyCDM predominately partition the nucleus into a single large bleb (i.e. Class I AsyCDM; ∼75%) and other classes of AsyCDM were less frequently observed (Figure 2D). In additional to NucRed staining, transfection with H2B-GFP also demonstrated the distribution of nuclear material into a single large bleb (Supplementary Figure 3B).

**Figure 2.**
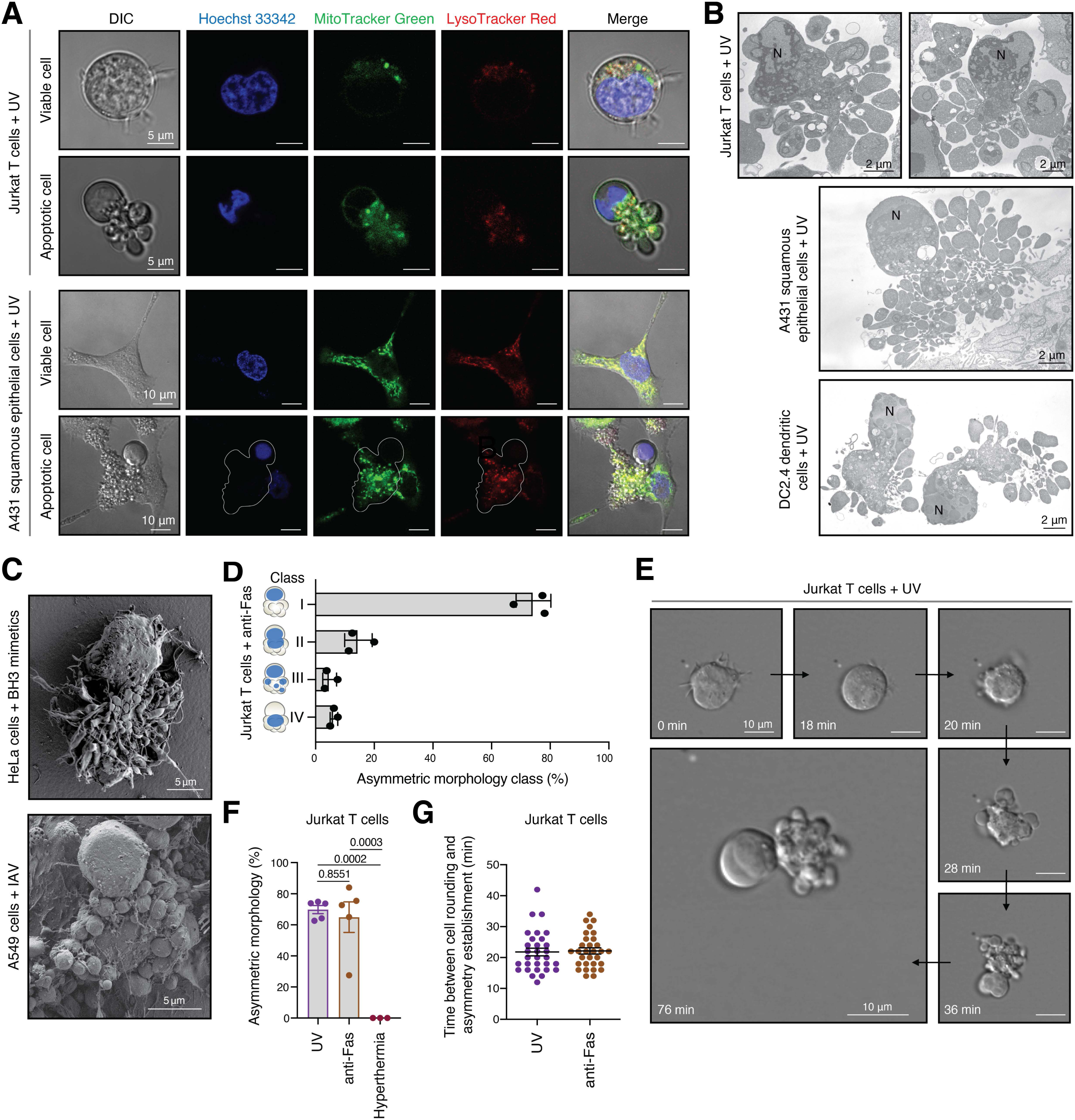
Establishment of asymmetric morphology during apoptosis *in vitro*. **(A)** Confocal microscopy images of viable and apoptotic Jurkat T cells stained with Hoechst 33342, MitoTracker Green, and LysoTracker Red. Cells were induced to undergo apoptosis by UV irradiation (150 mJ/cm^2^) and images were captured at 2-3 hrs post UV treatment. **(B)** Transmission electron microscopy images of apoptotic Jurkat T cells, A431 squamous epithelial cells and DC2.4 dendritic cells with asymmetric morphology. N, nucleus (with condensed chromatin). **(C)** Scanning electron microscopy images of apoptotic HeLa cells and A549 cells treated with BH3 mimetics (2.5 μM ABT-737/0.5 μM S63845) or infected with influenza A virus (IAV), respectively. **(D)** Quantification of the classification of nuclear position during the establishment of asymmetric morphology. Asymmetric morphology class percentage was determined as the proportion of cells in each class to the total number of cells that exhibit asymmetric morphology, as monitored by time-lapse confocal microscopy (Supplementary Figure 3A). (n>45, N = 3) Error bars represent s.e.m. **(E)** Cellular morphology of Jurkat T cells undergoing UV-induced apoptosis was monitored for 4 h by time-lapse DIC microscopy. The establishment of asymmetric morphology is defined by the presence of a clear single large bleb at one extremity and multiple dynamic blebs located at the opposite side of the apoptotic cell. **(F)** Apoptotic cell morphology was quantified by time-lapse DIC microscopy analysis over 4 hrs to determine the percentage of cells undergoing apoptosis that establish asymmetric morphology. (n>50, N = 5) Error bars represent s.e.m. **(G)** The kinetics of asymmetric morphology establishment during apoptosis (UV or anti-Fas treated Jurkat T cells) was analysed. (n>20, N = 3) Error bars represent s.e.m.

Next, to determine when AsyCDM would be established during the process of apoptotic cell disassembly, we monitored Jurkat T cells undergoing apoptosis by time-lapse DIC microscopy and noted that after the initiation of surface membrane blebbing, AsyCDM was established when the apoptotic cells were undergoing dynamic membrane blebbing (Figure 2E). AsyCDM was observed in the majority of Jurkat T cells treated with UV or anti-Fas to induce apoptosis (Figure 2F). It is worth noting that AsyCDM was not observed for Jurkat T cells undergoing primary necrosis induced by hyperthermic treatment (Figure 2F, Supplementary Figure 3C). The establishment of AsyCDM occurred at approximately 20 min after the cell rounding stage (i.e. the first morphological change during apoptosis for Jurkat T cells^15,18^) and was comparable between different apoptotic stimulus (Figure 2G). AsyCDM was also observed in a variety of cell types including mouse primary splenic monocytes and thymocytes undergoing UV- or dex-induced apoptosis, respectively (Supplementary Figure 3D). In comparison to apoptotic Jurkat T cells that don’t regularly undergo complete cell disassembly into ApoBDs^15,18^, DC2.4 (DC-like) cells frequently form ApoBDs during apoptosis (Figure 3A-B). Notably, for DC2.4 cells, we frequently (>80%) observed the establishment of AsyCDM during apoptosis and subsequent formation of smaller ApoBDs from the side of the apoptotic cell opposite to the largest bleb (Figure 3A-C). Since DC2.4 cells were able to generate an abundant of ApoBDs during apoptosis through the formation of thin membrane protrusions known as apoptopodia and beaded-apoptopodia^15,16^ (Figure 3A), we monitored DC-derived ApoBDs and found that the majority of ApoBDs lack DNA content and those that harbours DNA are larger (Figure 3C-E). The establishment of AsyCDM and subsequent release of a single large ApoBD from the dying cell was also observed for BH3 mimetics-treated DC2.4 and mouse bone marrow-derived DCs, as well as UV-treated human monocyte-derived DCs (HMDCs) (Figure 3F-H). Together, these data suggest that the establishment of AsyCDM are frequently observed in many cell types, which could aid the partitioning of nuclear material into one large bleb and subsequent packaging into the largest ApoBDs following apoptotic cell disassembly.

**Figure 3.**
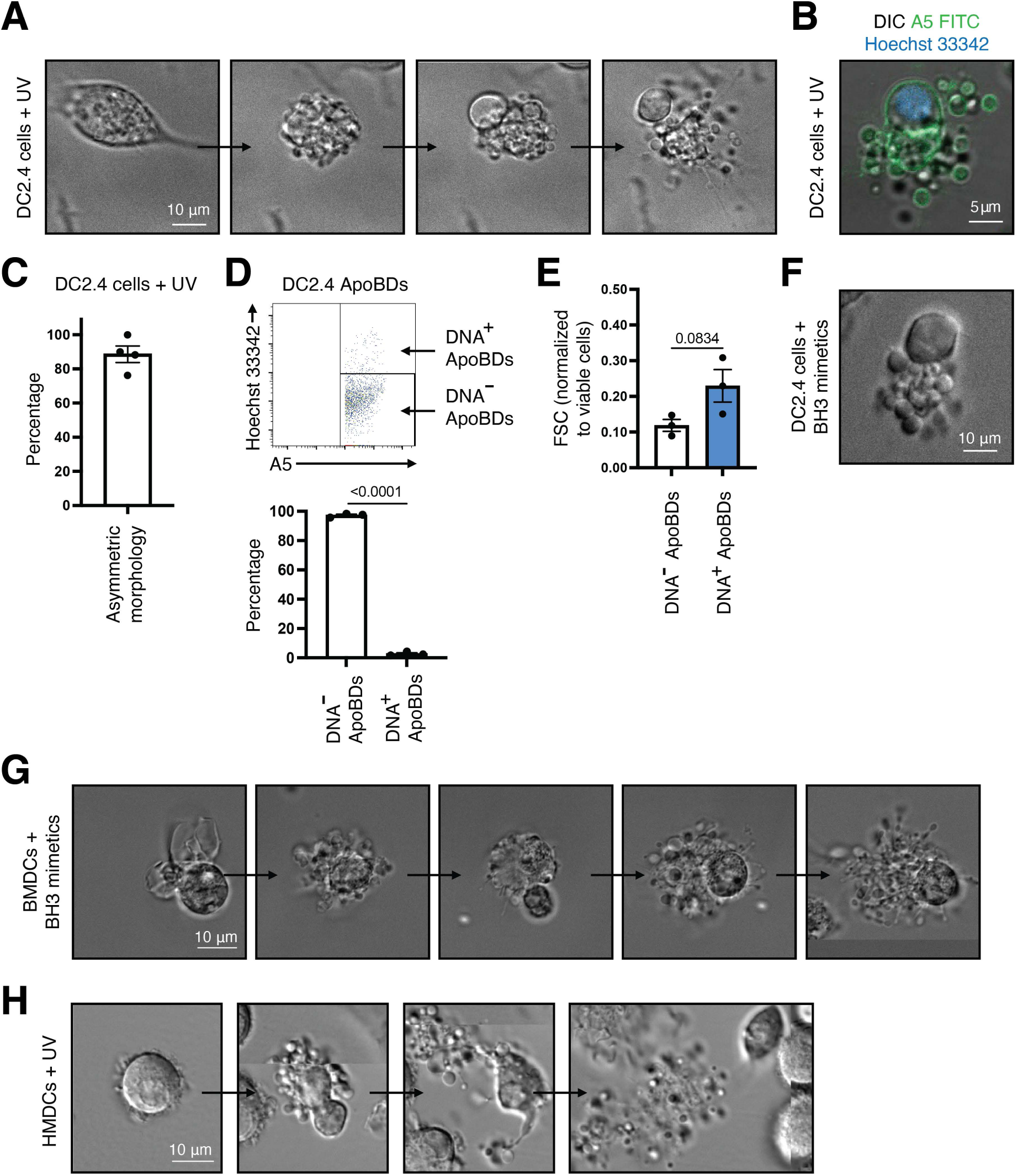
Establishment of AsyCDM and ApoBD formation by DCs. **(A)** Time-lapse DIC microscopy images monitoring the formation of membrane blebs and apoptopodia by DC2.4 cells treated with UV (150 mJ/cm^2^) to induce apoptosis. **(B)** A5-FITC and Hoechst 33342 staining of PtdSer and nuclear DNA, respectively, on/in DC2.4 cell induced to undergo apoptosis by UV (150 mJ/cm^2^). **(C)** Quantitation of live microscopy data from (A) to determine the percentage of apoptotic DC2.4 cells that underwent surface blebbing, dynamic blebbing, and apoptopodia formation. (*n* = 4) Error bars represent s.e.m. **(D)** Flow cytometry analysis of the distribution of DNA into ApoBDs generated from UV-treated DC2.4 cells. (*n* = 3) Error bars represent s.e.m. **(E)** Relative size of DNA^-^ and DNA^+^ ApoBDs generated from DC2.4 cells compared to viable cells following UV treatment. (*n* = 3) Error bars represent s.e.m. **(F)** DIC microscopy images monitoring the establishment of AsyCDM of DC2.4 cells induced to undergo apoptosis by BH3 mimetics. Data representative of (*n* = 3). Time-lapse DIC microscopy images monitoring the disassembly of UV-treated apoptotic BMDCs **(G)** and UV-treated apoptotic HMDCs **(H)**. Unpaired student’s two tailed *t*-test was performed to determine the indicated *p*-value.

### Role of erythroblast enucleation and cell polarity regulators in AsyCDM establishment

Partitioning of the nucleus to one extremity of apoptotic cells through AsyCDM appeared to be remarkably similar to erythroblasts undergoing enucleation, whereby the nucleus of an erythroblast is expulsed from the cell prior to the formation of mature red blood cell (Figure 4A-B). Similar to DNA-containing ApoBDs, the expulsed cellular fragment from erythroblast known as pyrenocyte harbours condensed chromatin and exposes the phospholipid phosphatidylserine (PtdSer) to aid phagocytic removal^4,27^. Comparable to erythroblastic islands whereby a central macrophage could aid erythropoiesis as well as the removal of pyrenocytes^28^, the presence of macrophages harbouring numerous apoptotic nuclei (known as tingible body macrophages) was also observed in the mouse thymus following dex-induced apoptosis (Figure 4C). Due to the parallels between these two processes, we next reason whether similar mechanisms are utilised by apoptotic cells to establish AsyCDM. We tested pharmacological compounds that are known to block erythroblast enucleation by inhibiting processes such as calcium signalling and vesicle trafficking. Treating apoptotic Jurkat T cells with EGTA (calcium chelator^29^), KN-96 (calmodulin-dependent kinase II (CaMKII) inhibitor^29^), and monensin (Na^+^ ionophore that impairs trafficking between endosomes and lysosomes^30^) had no significant effect on the proportion of blebbing cells undergoing AsyCDM (Figure 4D), suggesting that calcium signalling and vesicle trafficking were not involved in the establishment of AsyCDM.

**Figure 4.**
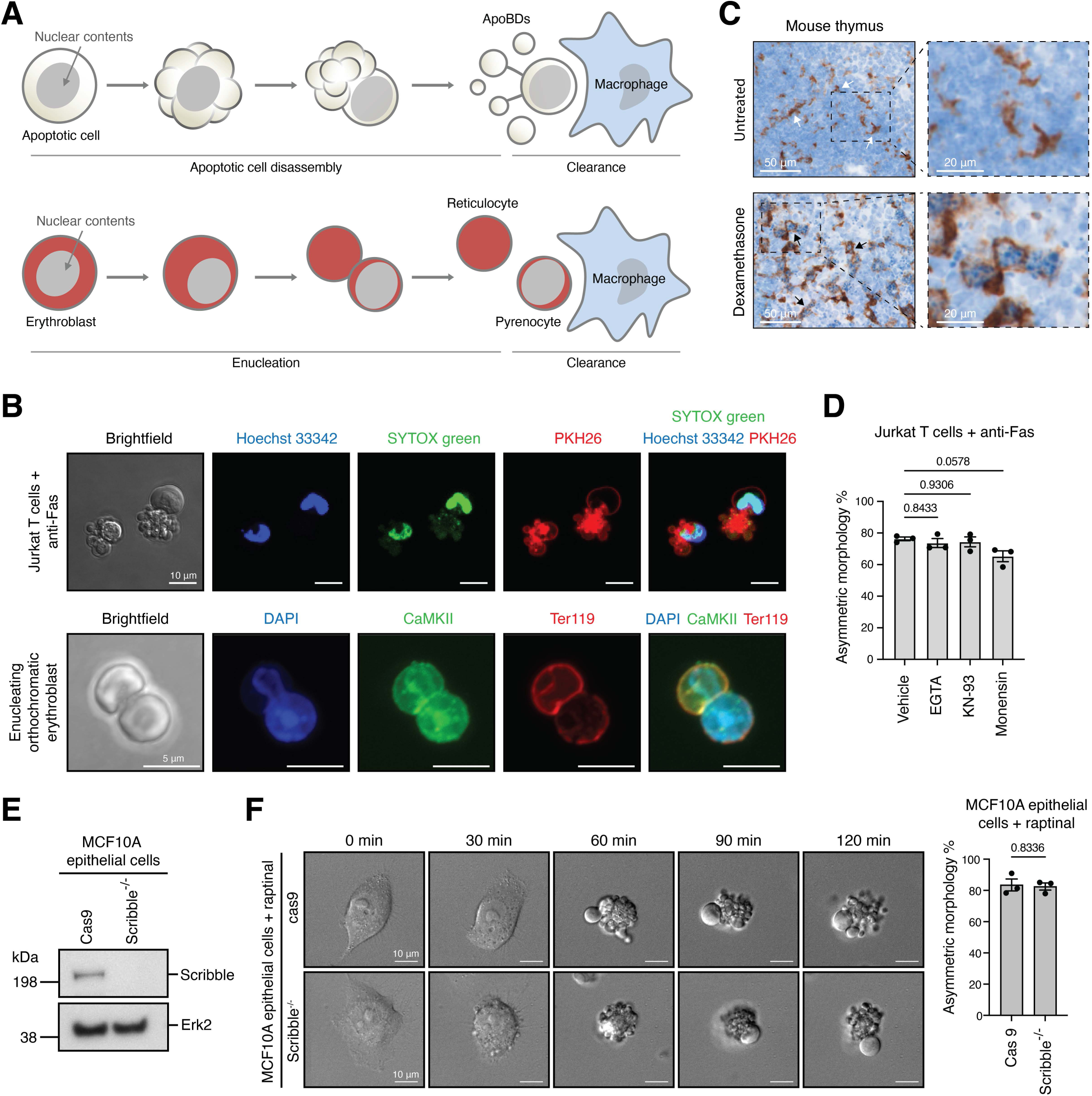
Role of RBC enucleation and cell polarity regulators in AsyCDM establishment. **(A)** Schematic of the apoptotic cell disassembly process in comparison to erythrocyte enucleation. **(B)** Confocal microscopy images of apoptotic Jurkat T cells treated with anti-Fas (250 ng/mL) and enucleating orthohromatic erythroblast. Cells were stained with Hoechst 33342, SYTOX green, PKH26, DAPI, CaMKII and Ter119 as indicated. Data representative of (*n* = 3). **(C)** Histological analysis of F4/80^+^ macrophages (brown, arrows) in the mouse thymus with and without dex-induced apoptosis (12.5 mg/kg). Nuclei (blue) are stained with Mayer Hematoxylin. **(D)** Quantification of Jurkat T cells undergoing anti-Fas (250 ng/mL) induced apoptosis treated with RBC enucleation inhibitors EGTA (1 mM), KN-93 (40 μM) and monensin (10 μM). Asymmetric morphology percentage was calculated as the proportion of cells that exhibit asymmetric morphology to the total number of cells undergoing dynamic plasma membrane blebbing, as monitored by time-lapse DIC microscopy. (n>45, N=3) Error bars represent s.e.m. **(E)** Immunoblot analysis of Cas9 and Scribble^-/-^ MCF10A epithelial cells for the expression of scribble. **(F)** Representative confocal time-lapse microscopy images of Cas9 and Scribble^-/-^ cells treated with 5 μM raptinal to induce apoptosis. Asymmetric morphology percentage was determined as per (D). (*n* = 3) Error bars represent s.e.m. One-way ANOVA followed by Dunnett test was performed for (D) and unpaired student’s two tailed *t*-test was performed for (F) to determine the indicated *p*-value. Whole thymic tissue was harvested and fixed in 4% paraformaldehyde (PFA) for 24 h and processed using the Tissue-Tek VIP 6 AI Tissue Processor (Sakura Finetek). Paraffin-embedded sections were immunohistochemically stained with anti-F4/80 (1:1,000, WEHI in-house antibody) and counterstained with Mayer Hematoxylin using the automated Omnis EnVision G2 template (Dako, Glostrup) as previously described^21^.

Comparable to the observations in this study, asymmetric cell division (ACD) results in the generation of two unique daughter cells with different molecular compositions^31^. This process is essential for generating cellular diversity and is regulated by various cytoskeletal components, with the establishment of cell polarity driven by the Scribble complex^31,32^. To determine if ACD regulator Scribble could contribute to the establishment of AsyCDM, we generated human mammary epithelial MCF10A cells lacking Scribble^19^, confirmed by immunoblot analysis (Figure 4E). Next, cells were treated with raptinal to induce apoptosis^33^ and cellular morphology was monitored by time-lapse DIC microscopy. Notably, lack of Scribble expression did not affect the establishment of AsyCDM during apoptosis (Figure 4F), suggesting that the key cell polarity regulator Scribble is not involved in the establishment of AsyCDM.

### Actomyosin contraction aids the establishment of AsyCDM

We next examine whether regulators of the morphological steps of apoptotic cell disassembly are involved in the establishment of AsyCDM by pharmacologic and genetic approaches (Figure 5A). First, dynamic membrane blebbing (i.e. Step 1 of apoptotic cell disassembly) is regulated by ROCK1^10,18,34^ and targeting ROCK1 using the small molecule inhibitor GSK 269962 markedly inhibited the establishment of AsyCDM by apoptotic Jurkat T cells (Figure 5A-C). Similarly, the establishment of AsyCDM was also reduced in ROCK1^-/-^ Jurkat T cells undergoing apoptosis as compared to Cas9 only control (Figure 5B and C). In contrast, targeting PANX1, a key negative regulator of apoptopodia formation (i.e. Step 2 of apoptotic cell disassembly)^15^, by using the PANX1 inhibitor trovafloxacin or PANX1^-/-^ Jurkat T cells did not affect the establishment of AsyCDM (Figure 5A-C). Collectively, these data suggest Step 1 but not Step 2 of apoptotic cell disassembly is required for the establishment of AsyCDM.

**Figure 5.**
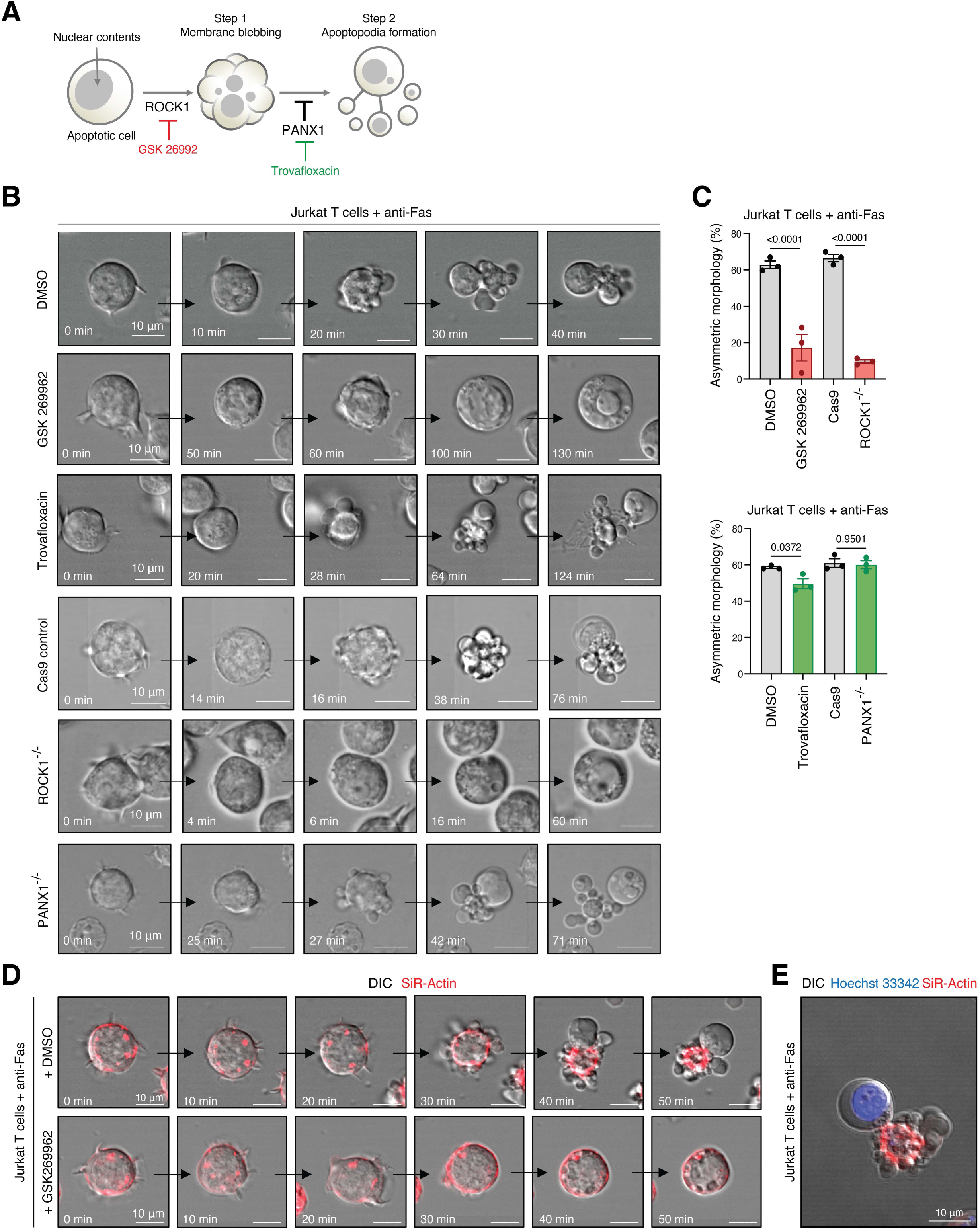
Role of apoptotic cell disassembly regulators on AsyCDM establishment. **(A)** Schematic of apoptotic cell disassembly and the molecular regulators of this process. **(B)** Confocal time-lapse microscopy images monitoring apoptotic morphology of DMSO, GSK 269962 or trovafloxacin treated Jurkat T cells, or Cas9 control, ROCK1^-/-^ or PANX1^-/-^ Jurkat cells. **(C)** Quantification of the percentage of Jurkat T cells as described in (B) that establish AsyCDM. Error bars represents s.e.m. (n>50, N=3). Cells were induced to undergo apoptosis by anti-Fas treatment (250 ng/mL). **(D)** Confocal time-lapse microscopy images monitoring the location of F-actin in Jurkat T cells undergoing anti-Fas (250 ng/mL) induced apoptosis in the presence or absence of the ROCK1 inhibitor GSK-269962. **(E)** Representative confocal microscopy image of Jurkat T cells undergoing anti-Fas (250 ng/mL) induced apoptosis stained with Hoechst 33342 and SiR-Actin. Data are representative of at least 3 independent experiments. Unpaired student’s two tailed *t*-test was performed to determine the indicated *p*-value.

It should be noted that the presence of small membrane blebs at one extremity of the apoptotic cell is a key-defining feature of AsyCDM (Figure 2) and since caspase-activated ROCK1 is essential in driving apoptotic membrane bleb formation through promoting actomyosin contraction^10,34^, it is expected that targeting ROCK1 could interfere with AsyCDM establishment (Figure 5B and C). Nevertheless, the importance of ROCK1 suggest that the contractile force generated downstream of ROCK1 activation may be required for AsyCDM establishment and partitioning of nuclear material into ApoBDs. Thus, we investigated the formation of F-actin filaments during the progression of Jurkat T cell apoptosis using a cell-permeable F-actin-binding probe SiR-Actin. F-actin was first observed in viable Jurkat T cells near the cell periphery and also in patches surrounding the cell nucleus prior to any apoptotic morphologies (Figure 5D). However, upon apoptosis progression, an F-actin ring appeared during the surface blebbing stage and continued to condense during the dynamic blebbing stage (Figure 5D). The majority of F-actin was found surrounding the central cell body, and not inside either the small blebs or the large nuclear containing bleb (Figure 5D and E). In contrast, when ROCK1 was targeted by GSK 269962 in Jurkat T cells undergoing apoptosis, F-actin staining was found predominately on the cell periphery and exhibited no dynamic changes during the progression of apoptosis (Figure 5D). Thus, the establishment of AsyCDM is dependent on ROCK1-mediated actomyosin contraction.

## Discussion

Dynamic morphological changes during apoptosis is critical for the formation ApoBDs prior to cell clearance^13,35^. Similarly, during the maturation of mammalian red blood cells, erythrocytes undergo dynamic morphological changes to enable the expulsion of the nucleus and subsequent removal by macrophages^36^. As noted in this study, although the establishment of AsyCDM during apoptosis and the erythrocyte enucleation process exhibit morphological similarities, and the formation of ApoBDs and pyrenocytes involve similar molecular processes such as caspase activation^10,34,37^, vesicular trafficking^16,30^ and extracellular calcium influx^29,38^, targeting key processes required for erythrocyte enucleation did not interfere with AsyCDM establishment. Thus, distinct mechanisms are utilised by apoptotic cells and erythrocytes to partition nuclear material for clearance. Nevertheless, it is worth noting that comparable F-actin ring at the junction between the nucleus containing bleb and other parts of the apoptotic cell during the establishment of AsyCDM has also been observed between reticulocyte and pyrenocyte during enucleation^39,40^, suggesting cytoskeletal rearrangement is key for both processes. Furthermore, the formation of the ‘F-actin basket’ around the nucleus during AsyCDM establishment also resembles the accumulation of actin around the nucleus of apoptotic neuroepithelial cells in *Drosophila*, which aids the relocation of the nucleus from the basal to apical side prior to cell fragmentation^41^. Interestingly, the movement of nuclei in apoptotic neuroepithelial cells contributes to the generation of apico-basal forces required for neural tube folding^40,42^. Whether the formation of nuclei-containing ApoBD through the establishment of AsyCDM could also exert mechanical forces on neighbouring cells and influence morphogenesis and/or tissue homeostasis remains to be defined.

Rapid removal of unwanted cellular contents such as the nucleus and mitochondria is essential to prevent the release of DAMPs and trigger inflammation. For example, pyrenocytes (containing the nucleus released from erythrocytes) are cleared in the bone marrow by macrophages^28,43^. Similarly, the release of mitochondria-containing large extracellular vesicles called exophers by cardiomyocytes are rapidly removed by neighbouring cardiac tissue macrophages^44^. Notably, the identification of ApoBD containing nuclear material in steady state in the thymus as well as in various thymocyte apoptosis models suggest the formation of this ApoBD subset could aid the removal nuclear contents from apoptotic cells. Notably, DNA^+^ ApoBDs are larger in size and macrophages have been shown to be more efficient at engulfing larger particles compared to non-professional phagocytes such as fibroblasts^18^. Thus, similar to the clearance of pyrenocytes by macrophages in the bone marrow, macrophages in the thymus could be the key phagocytes in mediating the removal of DNA^+^ ApoBDs. Consistent with this notion, the presence of nuclear containing apoptotic material in thymic macrophages are well documented^23,45^. Moreover, in DNase II deficient mice, macrophages in the thymus showed the accumulation of undigested DNA, further suggesting that macrophages is critical in mediating the degradation of nuclear DNA from apoptotic thymocytes^46^.

Collectively, data presented in this study demonstrates a novel mechanism in regulating the distribution of cellular contents during the progression of apoptosis, with implication in generating ApoBDs containing distinct contents such as nuclear material to aid cell clearance and potentially intercellular communication.

## Author contributions

J.P.S., L.J. and I.K.H.P. designed the experiments with input from co-authors, in particular A.L.H., M.D.H., T.K.P., R.T., S.S., A.A.B., P.O.H. and G.K.A.-S. J.P.S. and I.K.H.P. wrote the manuscript. All figures were assembled by J.P.S., L.J., A.L.H., I.K.H.P., D.C.O., S.F.R. and T.K.P. S.K., I.Y., L.J., D.C.O. and S.F.P. performed TEM and SEM experiments. S.C. and L.N. performed microscope analysis on A431 cells and enucleating orthohromatic erythroblast by confocal microscopy, respectively. R.T. generated ROCK1^-/-^ and PANX1^-/-^ Jurkat T cells. R.S. generated MCF10A Cas9 and Scribble^-/-^ cells. J.P.S. and L.J. performed all other experiments.

## Acknowledgements

The authors would like to thank the La Trobe University Bioimaging Platform, La Trobe Animal Research and Teaching Facility and Walter and Eliza Hall Institute histology team for their technical supports, and Prof Sarah Russell for helpful discussion. This work was funded by the National Health and Medical Research Council (GNT1173662 to I.K.H.P.) and Australia Research Council (DP200100458 to I.K.H.P.).

## Supplementary Figure Legends

**Supplementary Figure 1.**
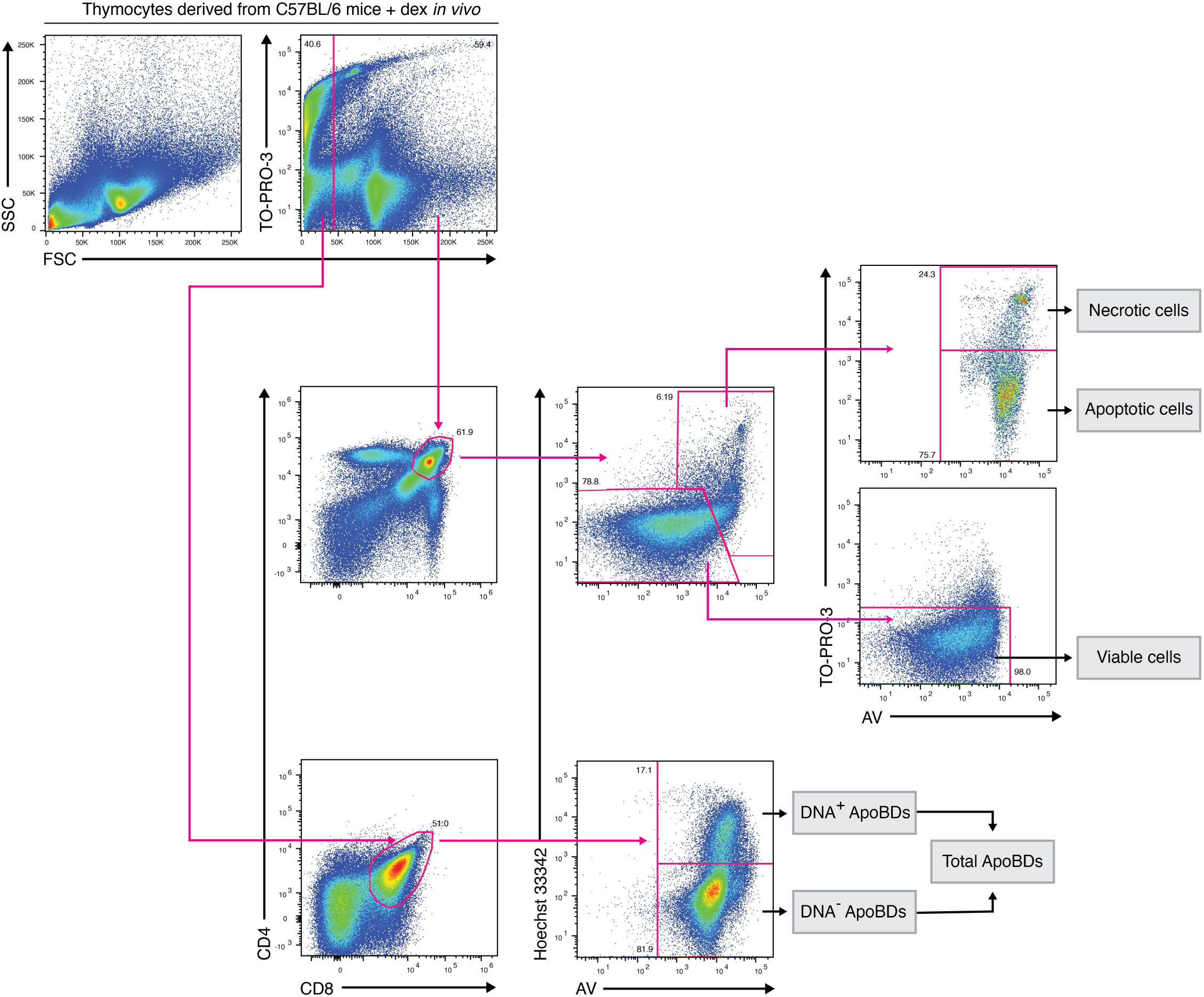
Electronic gating strategy for DNA^-^ and DNA^+^ thymocyte-derived ApoBDs. Flow cytometry analysis showing the electronic gating strategy used to identify thymocyte-derived ApoBDs that harbours high or low amount of DNA contents.

**Supplementary Figure 2.**
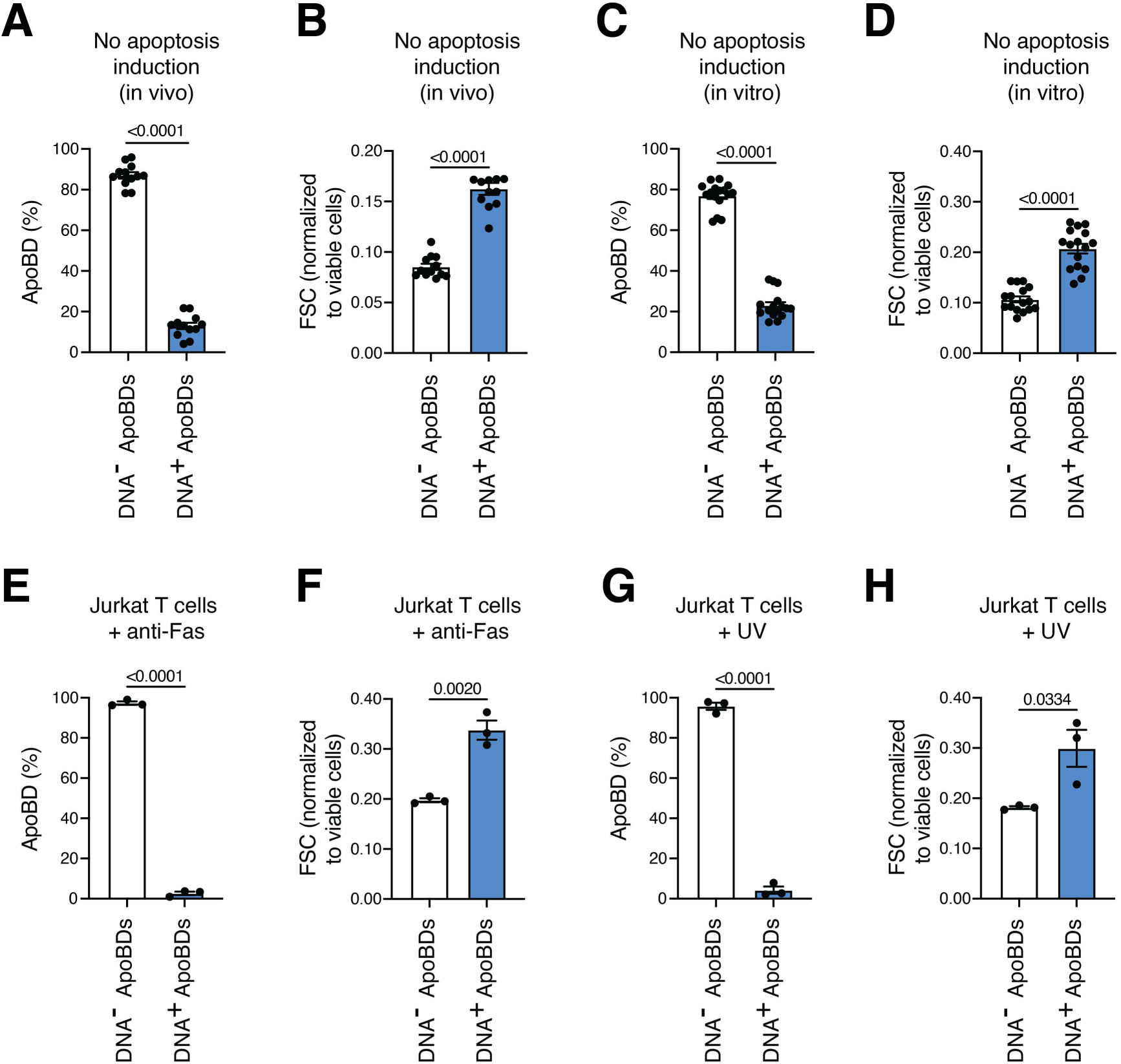
Formation of distinct DNA-containing ApoBDs in vivo and in vitro from mouse thymocytes and human Jurkat T cells. Flow cytometry analysis of the proportion of DNA^-^ and DNA^+^ ApoBDs derived from thymocytes **(A)** and their relative size compared to viable cells *in vivo* (B) in steady state. (*n* = 12) Error bars represent s.e.m. Flow cytometry analysis of the proportion of DNA^-^ and DNA^+^ ApoBDs derived from thymocytes **(C)** and the relative size compared to viable cells *ex vivo* (D) in the absence of additional apoptotic stimulus. (*n* = 16) Error bars represent s.e.m. **(E)** Flow cytometry analysis of the distribution of DNA into Jurkat T cell derived ApoBDs following treatment with 250 ng/mL anti-Fas. (*n* = 3) Error bars represent s.e.m. **(F)** Relative size of DNA^-^ and DNA^+^ ApoBDs derived from Jurkat T cells compared to viable cells following 250 ng/mL anti-Fas treatment. (*n* = 3) Error bars represent s.e.m. Flow cytometry analysis of the of the distribution of DNA into Jurkat T cell derived ApoBDs following treatment with UV irradiation 150 mJ/cm^2^ **(G)** and the relative size of DNA^-^ and DNA^+^ ApoBDs compared to viable cells (**H)**. Data representative of (*n* = 3). Error bars represent s.e.m. Unpaired student’s two tailed *t*-test was performed to determine the indicated *p*-value.

**Supplementary Figure 3.**
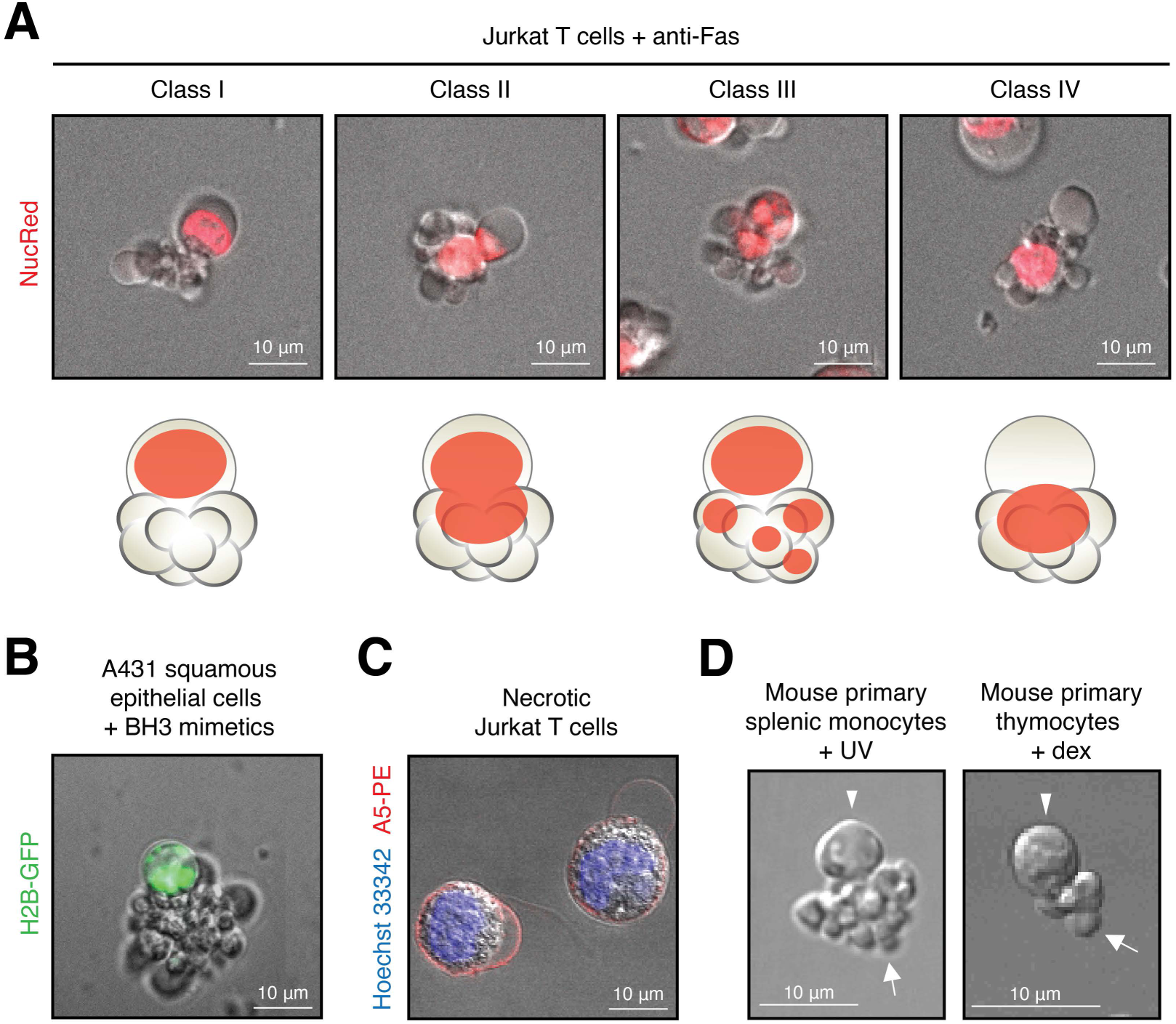
Characterisation of AsyCDM in different cell types and monitor the localisation of nuclear material. **(A)** Confocal microscopy images of Jurkat T cells undergoing anti-Fas (250 ng/mL) mediated apoptosis. Classification of AsyCDM based on the location of nuclear material as determined by NucRed staining. Data representative of (*n* = 3). **(B)** Confocal microscopy of H2B-GFP A431 squamous epithelial cells undergoing BH3 mimetic (2.5 μM ABT-737/0.5 μM S63845) mediated apoptosis. Data representative of (*n* = 3). **(C)** Confocal microscopy images of hyperthermic induced necrotic Jurkat T cells stained with Hoechst 33342 and A5-PE. Data representative of (*n* = 3). **(D)** Asymmetric cell death morphology was observed for UV-treated mouse primary splenic monocytes, and dex-treated mouse primary thymocytes. Data are representative of (*n* = 2).

